# Mechanical variations in proteins with large-scale motions highlight the formation of structural locks

**DOI:** 10.1101/221077

**Authors:** Sophie Sacquin-Mora

## Abstract

Protein function depends just as much on flexibility as on structure, and in numerous cases, a protein’s biological activity involves transitions that will impact both its conformation and its mechanical properties. Here, we use a coarse-grain approach to investigate the impact of structural changes on protein flexibility. More particularly, we focus our study on proteins presenting large-scale motions. We show how calculating directional force constants within residue pairs, and investigating their variation upon protein closure, can lead to the detection of a limited set of residues that form a *structural lock* in the protein’s closed conformation. This lock, which is composed of residues whose side-chains are tightly interacting, highlights a new class of residues that are important for protein function by stabilizing the closed structure, and that cannot be detected using earlier tools like local rigidity profiles or distance variations maps, or alternative bioinformatics approaches, such as coevolution scores.

---

A preliminary version of this work, https://doi.org/10.1101/221077, was deposited in bioRxiv.

## 1. Introduction

Decades of protein studies have now made it clear that flexibility is just as important as structure for defining a protein’s activity (Henzler-Wildman and Kern, 2007; Micheletti, 2013; Orozco, 2014; Teilum et al., 2009). Indeed, conformational diversity is an essential feature in a functional protein, whether it is involved in catalysis, signal transduction (Nussinov and Ma, 2012), recognition (Boehr et al., 2009), or allostery (Gunasekaran et al., 2004). Protein motions are difficult to observe directly using experimental approaches, as they can cover a wide range of timescales, from a few picoseconds to 10’s of millisecond or more (Dror et al., 2012). Therefore, numerous techniques are now available to study protein motions on a range of different timescales, such as NMR (Kovermann et al., 2016; Narayanan et al., 2017), single molecule approaches (Colomb and Sarkar, 2015), time-resolved X-ray crystallography (Levantino et al., 2015; Meisburger et al., 2017), FRET (Dimura et al., 2016; Lerner et al., 2018) or SAXS (Kikhney and Svergun, 2015), which can be used alone or combined together (Debiec et al., 2018). From a theoretical perspective, Molecular Dynamics (MD) simulations represent a classic alternative and will often complement experimental work (Debiec et al., 2018; Feng et al., 2016; Narayanan et al., 2017). Still, the use of all-atom MD remains a costly strategy to access rare events, such as slow, large-amplitude conformational transitions, and usually requires the development of enhanced sampling strategies (Greener et al., 2017; Maximova et al., 2016; Romanowska et al., 2012; Seyler and Beckstein, 2014), or access to remarkable computational power (Shaw et al., 2010). Coarse-grain models, based on simplified protein representation and energy functions, are a second option to investigate structural changes that are not accessible to all-atom MD simulations (Al-Bluwi et al., 2013; Poma et al., 2017; Tiwari and Reuter, 2017; Zheng and Wen, 2017). Elastic Network Models (ENM) combined with Normal Mode Analysis (NMA) in particular, have proven particularly useful in this regard, since a small number of low-frequency normal modes will provide information regarding large-amplitude conformational changes in a protein (Al-Bluwi et al., 2013; Fuglebakk et al., 2015; Kurkcuoglu et al., 2016; Lopez-Blanco and Chacon, 2016; Mahajan and Sanejouand, 2015; Tama and Sanejouand, 2001; Uyar et al., 2014; Yang et al., 2007).

In that perspective, the ProPHet (for Probing Protein Heterogeneity) program, which combines a coarse-grain ENM with Brownian Dynamics (BD) simulations, was initially developed to investigate protein mechanical properties on the residue level (Sacquin-Mora and Lavery, 2006). Later work on proteins undergoing functionally related structural transitions (such as enzymes) showed that the mechanical variations induced by these conformational changes target a limited number of specific residues occupying key locations for enzymatic activity (Sacquin-Mora, 2016). In a previous, multi-scale, study on the conformational transition of guanylate kinase, we showed that the enzyme closure leads to the formation of a *structural lock* constituted of residues forming a dense set of interactions and stabilizing the protein’s closed form (Delalande et al., 2011). In this study, we used ProPHet to investigate the mechanical variations associated with structural changes for a set of 53 proteins, 13 of which exhibit large-scale motions (LSM) leading to root mean-square deviations (RMSDs) > 4 Å. Calculations of directional force constant variations within residue pairs show the formation of similar structural locks in the closed structures of 12 out of 13 proteins. Interestingly, neither residue specific force constants, nor distance variation maps or normal mode analysis permit to detect these functionally relevant residues. Additional calculations of coevolution scores for residue pairs also fail to predict the location of the structural lock within the tested proteins.

## 2. Material and Methods

**Coarse-grain Simulation:** Coarse-grain Brownian Dynamics (BD) simulations were performed using the ProPHet program (http://bioserv.rpbs.univ-paris-diderot.fr/services/ProPHet/) (Lavery and Sacquin-Mora, 2007; Sacquin-Mora, 2014; Sacquin-Mora et al., 2007a) on a set of 53 proteins listed in table SI-1 in the Supporting Material, for which the unbound/ligand-bound or open/closed structures are available in the Protein Data Bank (PDB). Note that 31 of these proteins are enzymes that were already investigated in more details in ref. (Sacquin-Mora, 2016). The root mean square deviations (RMSDs) between the unbound and ligand-bound (respectively the open and closed) structures ranges from 0.17 Å (for endo-xylanase) to 15.6 Å (for diphteria toxin), and 13 proteins are considered to exhibit large-scale conformational transitions, with RMSDs > 4 Å. While early CG-models would describe each residue by a single pseudo-atom (Tozzini, 2005), ProPHet uses a more detailed model enabling different residues to be distinguished, in a similar fashion as more recent CG protein representation (Kmiecik et al., 2016) like the now classic MARTINI model (Marrink and Tieleman, 2013), OPEP (Sterpone et al., 2014) or PaLaCE (Pasi et al., 2013). The amino acids are represented by one pseudo-atom located at the C*α* position, and either one or two (for larger residues) pseudo-atoms replacing the side-chain (with the exception of Gly) (Zacharias, 2003). Interactions between the pseudo-atoms are treated according to the standard Elastic Network Model (ENM) (Tozzini, 2005), that is, pseudoatoms closer than the cutoff parameter, *Rc* = 9 Å, are joined by gaussian springs that all have identical spring constants of *γ* = 0.42 N m^−1^ (0.6 kcal mol^−1^ Å^−2^). The springs are taken to be relaxed for the starting conformation of the protein, i. e. its crystallographic structure. Note that, since the seminal works of Tirion (Tirion, 1996) and Bahar et al. (Bahar et al., 1997), numerous variations of elastic networks models have been proposed (Fuglebakk et al., 2015). Refinements include distance dependent force constants (Hinsen, 1998), specific force constants for nearest neighbor pairs (Hinsen et al., 2005), different parameters to model inter and intra domain contacts (Song and Jernigan, 2006), or parameter enabling the breaking of contacts during the simulation in order to model conformational transitions (Putz and Brock, 2017; Zheng and Wen, 2017). However, comparative studies have shown that from a qualitative point of view, elastic network models appear to be remarkably robust (Leioatts et al., 2012), and since the purpose of this work is to model proteins mechanics in their equilibrium state, and not the conformational transition between two structures, we used the original, single-parameter, model.

The simulations used an implicit solvent representation via the diffusion and random displacement term in the equation of motion (Ermak and McCammon, 1978),

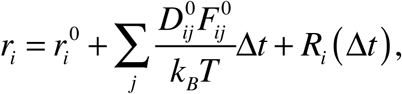

where **r**_i_ and **r**_i_^0^ denote the position vector of particle *i* before and after the time step *Δt*, **F**_i_ is the force on particle *i*, **R**_i_(*Δt*) is a random displacement, and hydrodynamic interactions are included through the configuration-dependent diffusion tensor **D** (Pastor et al., 1988). We used a bulk solvent viscosity η = 1.0 cP (1 P being 0.1 Pa.s), corresponding to water at room temperature. Mechanical properties are obtained from 200,000 BD steps at 300 K. The simulations are analyzed in terms of the fluctuations of the mean distance between each pseudo-atom belonging to a given amino acid and the pseudo-atoms belonging to the remaining protein residues. The inverse of these fluctuations yields an effective force constant *k_i_* that describes the ease of moving a pseudo-atom *i* with respect to the overall protein structure:

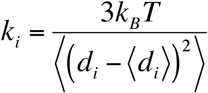

where〈〉 denotes an average taken over the whole simulation and *d_i_* =〈*d_ij_*〉*_j*_* is the average distance from particle *i* to the other particles j in the protein (the sum over *j\** implies the exclusion of the pseudo-atoms belonging to the same residue as i). The distances between the C*α* pseudo-atom of residue *k* and the C*α* pseudo-atoms of the adjacent residues *k-1* and *k+1* are excluded, since the corresponding distances are virtually constant. The force constant for each residue *k* in the protein is the average of the force constants for all its constituent pseudo-atoms *i*. We will use the term *rigidity profile* to describe the ordered set of force constants for all the residues in a given protein. Note that, following earlier studies which showed how small ligands had little influence on calculated force constants (Sacquin-Mora and Lavery, 2006; Sacquin-Mora et al., 2007a), we chose not to include ligands in the protein representations. This enables us to study the proteins intrinsic mechanical properties independently of the nature and position of any bound ligand.

In ProPHet, we can also use the fluctuations of the inter-residue distances *d_ij_* to compute force constants similar to those obtained by Eyal and Bahar (Eyal and Bahar, 2008) with a one-point-per-residue ENM. The resulting directional force constants (DFCs) are simply calculated as

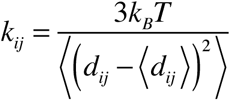
and can be used to construct a complete map of the mechanical resistance of a protein in response to all possible pulling directions.

**Calculation of coevolution scores:** Coevolution scores for residue pairs were calculated using the CoeViz tool (Baker and Porollo, 2016) on the Polyview web-server (http://polyview.cchmc.org/) (Porollo et al., 2004) using the default parameters. Given a target protein sequence, a multiple sequence alignment (MSA) based on the UniProt UniRef90 clusters (Suzek et al., 2015) is generated using three iterations of PSI-BLAST (Altschul et al., 1997) with 2000 aligned sequences. CoeViz computes coevolution scores from the MSA using three different covariance metrics: chi-square statistics, **χ**^2^ (Larson et al., 2000), mutual information MI (Clarke, 1995), and Pearson correlation (see equations 1–3 in ref. (Baker and Porollo, 2016)). The tool also computes one conservation metric, which is the joint Shannon entropy (JE), see equation 4 in ref. (Baker and Porollo, 2016).

## 3. Results and Discussion

The rigidity profiles and mechanical maps of the open/unbound and closed/ligand-bound structures of the proteins listed in table SI-1 were systematically compared in order to investigate the mechanical changes induced by conformational variations.

### General considerations

There is a general positive correlation between the amplitude of the conformational variation in a protein and its average mechanical variation, as can be seen on Figure 1a-b (with correlation coefficients of 0,80 and 0.65 for the average force constant variation 〈Δk〉 and the root mean square force constant variation RMSΔk, respectively). As shown in a previous work made specifically on proteins undergoing domain-domain motions (Sacquin-Mora, 2014), large hinge-bending-type conformational transitions involve the closure of one domain on the other, usually resulting in a larger buried surface in the interdomain interface and leading to a more rigid structure. This is for example the case of the T4 lysozyme mutant (see Figure 2a-c), where protein closure induced by hinge motion of the two domains leads to a large rigidity increase for catalytic reside Glu11, ligand binding residues Gly30 and Met102 (Merski and Shoichet, 2012; Shoichet et al., 1995; Weaver and Matthews, 1987), and channel lining residues Met106 and Trp138 (Collins et al., 2007). On the contrary, cavity lining residues Met6 and Cys97, which form hydrogen bonds with water molecules (Dixon et al., 1992), see a decrease in their force constant upon proteins closure, thus suggesting a disruption of this interaction network. More generally, one must keep in mind that the force constants variations resulting from conformational change are highly heterogeneous along the protein sequence and cannot simply be predicted from the RMSD between the initial and final structures. Early work on the bacterial reaction center showed that point mutations with negligible structural impact (RMSD between the wild-type and the mutant proteins below 0.1 Å) can still have a large impact on the protein local flexibility and function (Sacquin-Mora et al., 2007b). Here, conformational changes upon ligand binding with small RMSDs (under 0.6 Å) can both lead to minor mechanical changes (see the case of ribonuclease MC1 in Figure 2d-f), or major mechanical variations for a limited set of specific residues. This is the case of ligand bound galactose mutarotase, see Figure 2g-i, where hinge residues Asn175, Ser177 and Glu307 undergo an important rigidity loss, and additional examples can be found in Ref. (Sacquin-Mora, 2016). Moreover, a small average mechanical variation over the whole protein can also result from a mixed mechanical response to structural change, with some residues (usually located in the active site) becoming more rigid upon ligand binding, while others (frequently lying on hinges and domain interfaces) become more flexible (see the case of carbonic anhydrase in ref. (Sacquin-Mora, 2016)).

**Figure 1.**
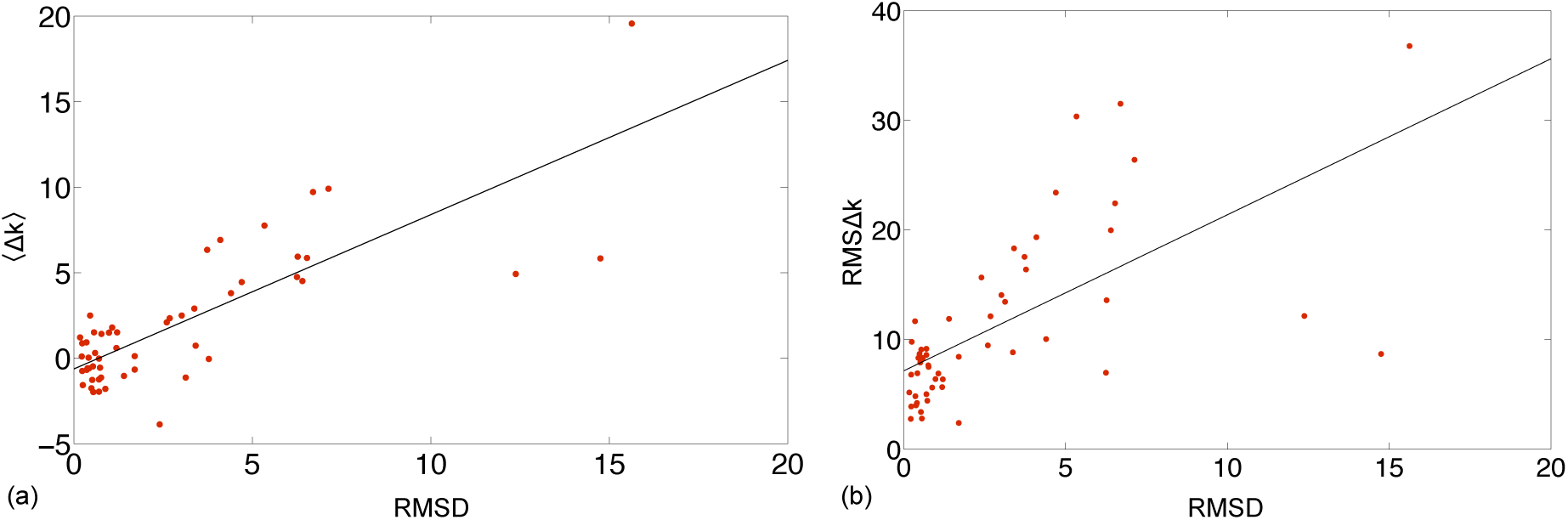
(a) Average force constant variation (in kcal mol^−1^ Å^−2^) upon closure/ligand binding as a function of the RMSD for the 53 proteins in the dataset. (b) Root mean square force constant variation (in kcal mol^−1^ Å^−2^) as a function of the RMSD for the 53 proteins in the dataset. The solid black lines show the best linear fit to the data.

**Figure 2.**
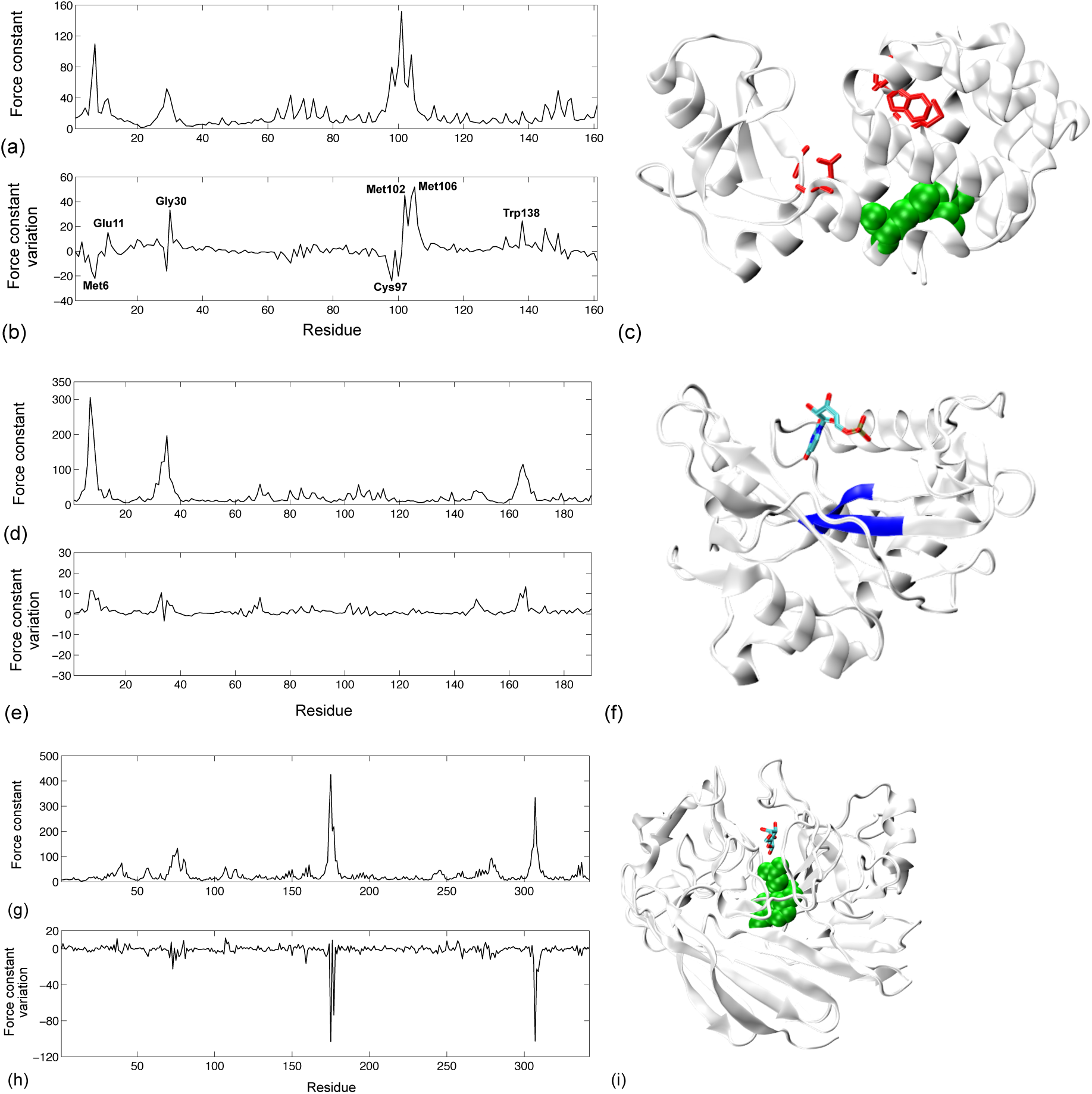
(a) Rigidity profile for T4 lysozyme mutant in its open form (all the rigidity curves are in kcal mol^−1^ Å^−2^) (b) Force constant variations upon closure (c) Cartoon representation of T4 lysozyme mutant in its closed form. Residues undergoing a large increase of their force constant upon closure are shown as red sticks, and residues undergoing an important rigidity decrease upon closure are shown as green van der Waals spheres. This color code is valid for all figures in the manuscript and supporting materials. Figures 2, 3, 6, 8 and figures in the supporting material were prepared using VMD (Humphrey et al., 1996). (d) Rigidity profile for ribonuclease MC1 in its unbound form (e) Force constant variations upon ligand binding (f) Cartoon representation of ribonuclease MC1 in its ligand bound form with the rigid protein core shown in blue. (g) Rigidity profile for galactose mutarotase in its unbound form (h) Force constant variations upon ligand binding (i) Cartoon representation of galactose mutarotase in its ligand bound form.

Another example of a mixed mechanical response can be seen for the N-terminal lobe from human serum transferrin (HST) in Figure 3. HST binds ferric ions in the bloodstream and transports them to cells. Its N-lobe (residues 1–337 of the native protein) folds in two domains (N1, residues 1–93+247–315, and N2, residues 94–246) separated by a hinge region. Upon ligand biding, the protein presents a rigid body rotation of the N2 domain relative to the N1 domain leading to a RMSD of 6.7 Å, while the RMSD calculated over individual domain are below 0.6 Å. This closure motion leads to an important increase in the rigidity of a limited set of residues (see Figure 3b), that are all involved either in the iron binding site, Asp63, Tyr188, His249 and Lys296, or the carbonate anion binding site, Thr120 and Arg124 (see Figure 3c). Meanwhile, hinge residues Ala244, Val246 and Pro247, which lie on the domain-domain interface, become more flexible upon closure.

**Figure 3.**
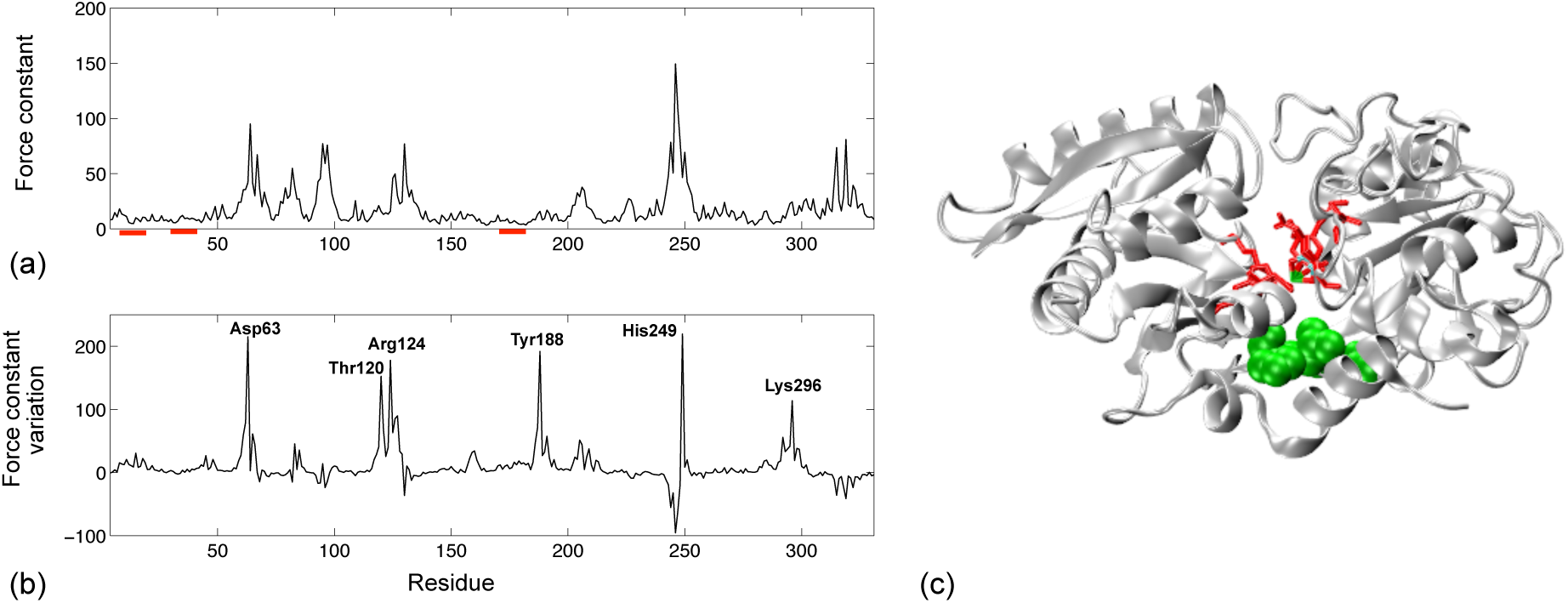
(a) Rigidity profile for human serum transferrin in its open form. The red bars highlight the residues involved in a structural lock after closure (b) Force constant variations upon closure (c) Cartoon representation of human serum transferrin in its closed form, the ionic Fe^3+^ ligand is shown as a green dot in the center.

Interestingly, we can also observe a negative correlation (with a correlation coefficient of −0.78) between the proteins average rigidity and the RMSD between their open/unbound and closed/ligand-bound structures (see Figure 4). Proteins that are, on average, more flexible, are also more likely to undergo large conformational transitions. More specifically, if we define *soft* proteins as those with an average rigidity below 20 kcal mol^−1^Å^−2^, 10 out of 13 elements from the large-scale motion group can be considered as soft proteins, while only 3 out of the 40 remaining proteins in the dataset belong to the same category. A noticeable outlier is the trp-repressor from E. coli, a small protein that appears to be remarkably flexible (with 〈k〉 = 4.7 kcal mol^−1^ Å^−2^) despite the small RMSD (1.70 Å) between its open and closed forms. However, after searching the PDB for additional conformations of this protein, we found an NMR structure of the trp-repressor complexed with DNA (PDB entry 1rcs). The RMSDs between the 15 conformers for this DNA-bound form and our reference open form of the trp-repressor are comprised between 5 and 5.6 Å (shown by a green point in Figure 4), so eventually the low average flexibility calculated for this protein accurately reflects its ability to undergo large conformational changes. The second point lying remarkably low under the black line in Figure 4 corresponds to importin β, which we expected to display a larger RMSD between its open and closed structures considering its weak rigidity (〈k〉 = 7.3 kcal mol^−1^ Å^−2^) in the open form. Once again, searching the PDB leads to the identification of a complexed form of importin β (bound to snuportin 1, PDB entry 2q5d), that displays a 7.7 Å RMSD from the original open structure (second green point in Figure 4). Furthermore, using streching molecular dynamics simulations, Kappel et al. (Kappel et al., 2010) also showed how importin β displays remarkable flexibility, thus agreeing with the coarse-grain simulations in the present study. The inclusion of these two corrected values increases the correlation coefficient between RMSD and 〈k〉 to 0.84. In addition, this correlation between 〈k〉 and the RMSD means that one should be able to estimate the amplitude of the conformational change undergone by a protein based on its general mechanical properties and without having any prior information regarding its final state.

**Figure 4.**
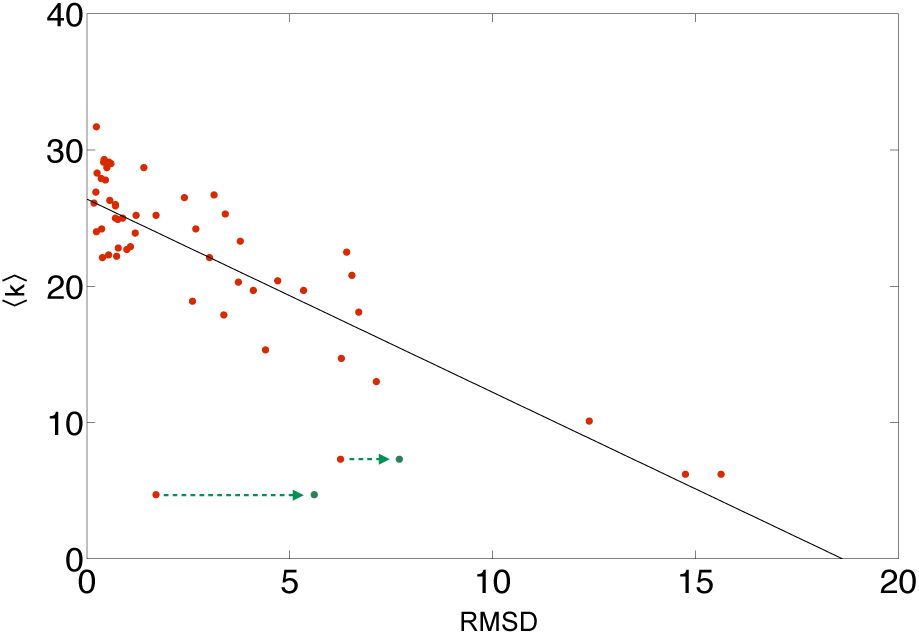
Average force constant (in kcal mol^−1^ Å^−2^) as a function of the RMSD (in Å) for the 53 proteins in the dataset in their open/unbound form. The solid black line shows the best linear fit to the data.

### Proteins undergoing large conformational transitions

**Variations in the mechanical map:** For the 13 proteins presenting large (with RMSDs > 4 Å) conformational transitions between their open/unbound and closed/ligand bound states, we also investigated the proteins DFC matrices and the variation in these mechanical maps resulting from the structural change.

DFC matrices for the proteins in their open/unbound state highlight the connections between pairs of residues belonging to the same protein domains, which will present higher directional force constants than pairs of residues belonging to different domains. For example, in Figure 5a one can clearly see how the high DFC areas correspond to the N1 and N2 domains in human serum transferrin. After closure, we can observe a shift of the DFC distribution toward larger values (see the histograms in Figure 5c), with the disappearance of many residue pairs with low force constants (below 2 kcal mol^−1^ Å^−2^). The evolution of the mechanical map shown in Figure 5b indicates that the increase in DFC values upon protein closure is restricted to a small number of inter-domain residue pairs. By selecting the residue pairs that present the largest relative increase (i. e, with at least 80% of the maximum relative increase observed for all pairs) in their DFC induced by conformational change, three structural groups are detected (see Figure 6). These residues come in close contact upon protein closure, which might explain the strong increase in the associated DFCs, with their side-chains tightly interacting, thus forming a *structural lock* that would help maintaining the protein in its closed conformation. Experimentally, the interactions between residue Val11 and Ser12 from the N1 domain, and Ser180 and Thr181 from the N2 domain involve hydrogen bonds through two buried, highly ordered water molecules (Jeffrey et al., 1998). In addition, residue Glu15 is highly conserved and forms hydrogen bonds with the opposite wall of the cleft it is lining (Haridas et al., 1995). More generally, all structural lock residues identified during this study are listed in table SI-2. Searching the literature for the led to the identification of several lock residues which had already been identified as functionally relevant in earlier experimental or theoretical studies, and which are highlighted in red in table SI-2. Most of these residues belong to conserved motives, and line the ligand binding pocket or the inter-domain interfaces, where they are likely to form salt-bridges or hydrogen bonds, while not being directly involved in the protein catalytic activity.

**Figure 5.**
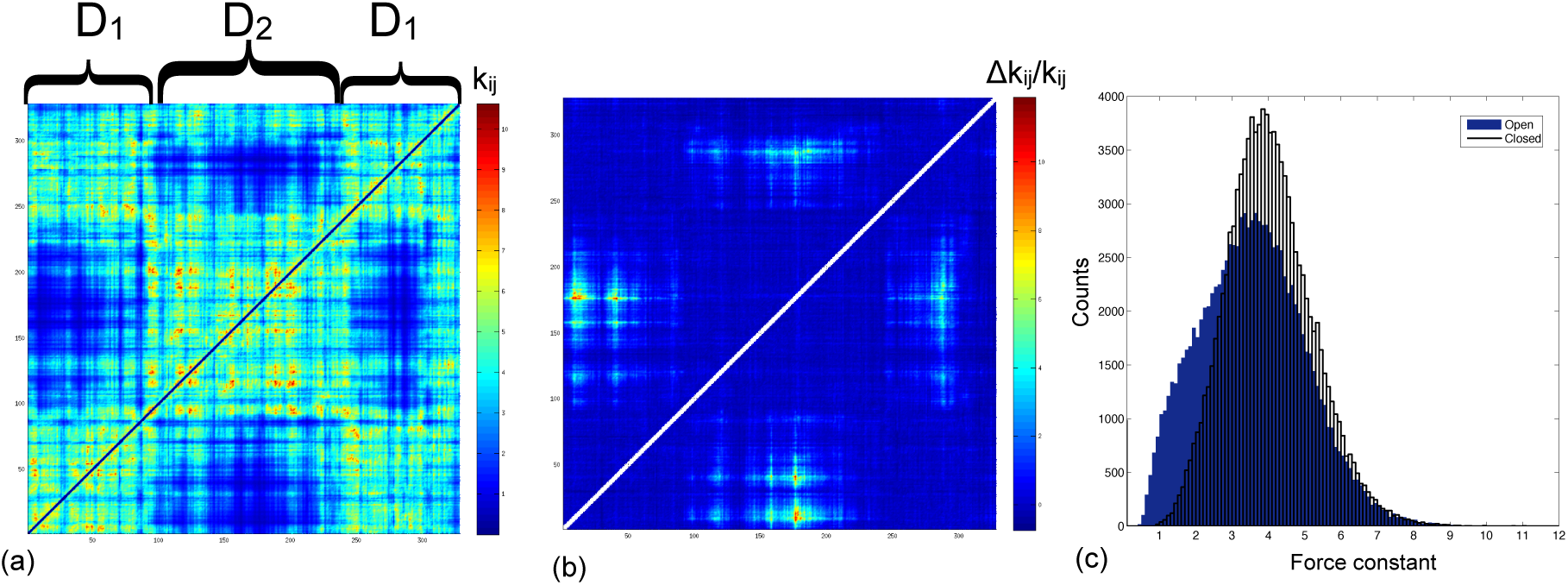
(a) Directional force constants matrix for human serum transferrin in its open form with the domains distribution along the sequence (b) Relative variation of the DFCs upon closure for human serum transferrin (c) DFCs distribution of human serum transferrin for the open (blue bars) and closed (transparent bars with black contours) conformations.

**Figure 6.**
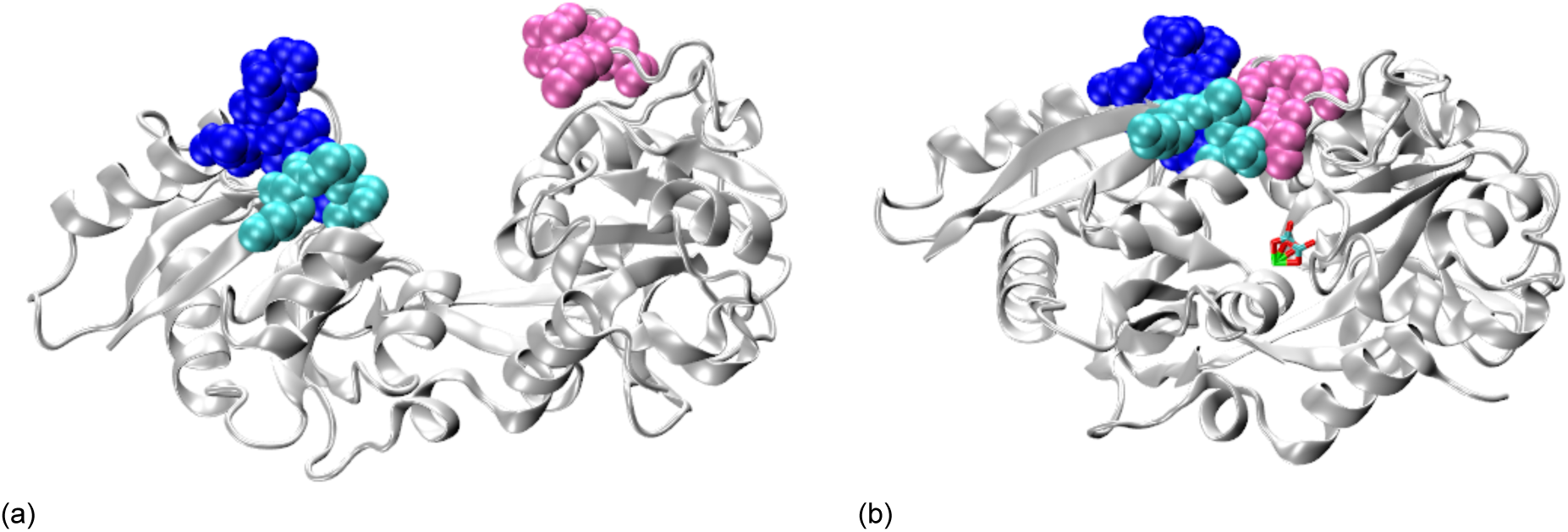
Cartoon representations of human serum transferrin with the residues forming rigid pairs upon closure shown as van der Waals spheres: Val11, Ser12, Glu13, His14, Glu15 and Ala16 (blue), Lys42, Ala43 and Ser44 (cyan), Cys179, Ser180, Thr181 and Asn183 (purple), (a) Open structure, (b) Closed structure.

Interestingly, when considered on their own, the residues involved in this structural lock are not rigid in the initial open protein conformation (their positions along the sequence are shown as red bars on Figure 3a), neither do they become rigid upon closure, see Figure 3b underneath. This means that components of the structural lock cannot be detected individually from simple rigidity profiles. Nevertheless, after closure they belong to rigid pairs, and their detection thus necessitates the use of mechanical maps that will highlight their functional role in the protein. Residue Trp50 from diphteria toxin (shown in dark blue in figures SI-16e and f) is an interesting example. It is located in a loop positioned over the NAD-binding pocket, and the W50A mutant of diphteria toxin displays a complete loss of its NADase activity (Wilson et al., 1994). Our coarse-grain calculations show that Trp50 is flexible in both the open and closed forms of diphteria toxin (see Figure SI-16a). To model the impact of the W50A mutation on the protein mechanical properties, we simply removed from the coarse-grain model the two pseudo-atoms corresponding to the tryptophan side-chain. As can be seen in Figures 7a-c this slight change in the elastic network has no visible effect on the closed protein rigidity profile and DFC maps. However, if we now consider the relative DFC variations upon closure, we can see in Figures 7d and e that they are much smaller in the W50A mutant compared to the native protein. While in the native diphteria toxin, residue pairs from the structural lock display a 70-fold increase in their DFC upon closure, in the W50A mutant, the maximum DFC increase lies below 18-fold. This result supports the hypothesis that the experimentally observed loss of catalytic activity for the W50A mutant might be related to a loss in stability for the protein in its closed form.

**Figure 7.**
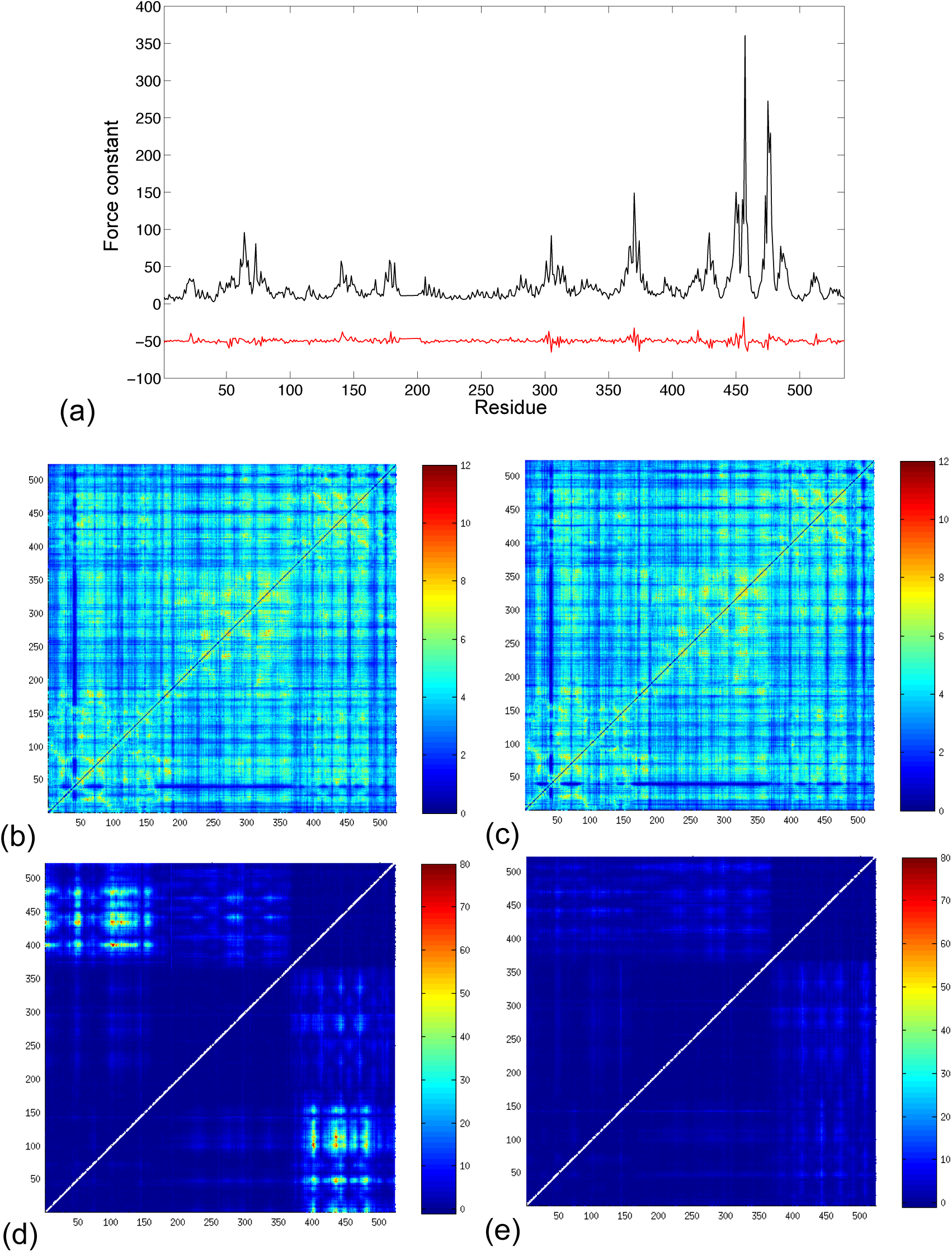
(a) Black line, rigidity profile for diphteria toxin in its closed form. Red line, variations in the force constant for the W50A mutant of diphteria toxin in its closed form compared to the native protein (with a −50 kcal mol^−1^ Å^−2^ vertical shift for visibility). DFC matrices for diphteria toxin in its closed form (with the same color scale 0:12): (b) Native protein, (c) W50A mutant. DFCs relative variation upon closure for diphteria toxin (with the same color scale −1:80): (d) Native protein, (e) W50 mutant.

DFC maps are qualitatively close to the mechanical resistance maps that were obtained by Eyal and Bahar by calculating effective force constants in response to uniaxial extensional forces exerted at each pair of residues (Eyal and Bahar, 2008). These also display the protein domains, and the variation in the mechanical resistance map upon protein closure highlights the same residue pairs, see on Figures SI-1b and c the maps calculated for human serum transferrin using the ProDy package (Bakan et al., 2014). However, the mean resistance profiles (and their variation upon protein closure) obtained from this approach, and which can be compared to our mechanical profiles, present much less contrast between flexible and rigid residues and do not permit to discriminate individual residues that are important for protein function (see Figure SI-1a). In addition, one can note that DFC maps are similar to the contact maps based on the inter-residue distances only (see for exemple the case of human serum transferrin in Figure S2–1a), with both matrices showing the protein domain composition along the sequence. DFC variations however, will highlight a much more limited set of residue pairs than simply looking at variation in the residue distances upon protein closure (cf. Figure S2–1b). The use of a binary contact maps also fails to account for the structural lock formation. In Figure SI-3a, the white points indicate residue pairs forming new contacts in the ENM (with a 9 Å cutoff) upon closure. The protein large-scale motion induces the formation of up to 14 new contacts for residues that are distributed all over the sequence (see figure SI-3b). If one focusses only on residues forming 10 or more new contacts upon closure, we can see in Figure SI-3c that they are preferentially located on the interface between the two protein domains, but do not coincide with the structural lock residues. Finally, the correlation matrix, displaying correlations between the motions of the Cα atoms in the structure over all the non-trivial normal modes, was calculated for human serum transferrin using the WEBnm@ server (Tiwari et al., 2014). Again, the variation in the correlated motions upon protein closure does not permit to highlight structural lock residues (see Figure SI-4). Altogether, residues forming the protein structural lock are not only residues getting nearer, or forming new contacts in the protein closed form. They present specific mechanical properties, that are related to the global protein dynamics, and cannot be identified using structural information only.

Similar structural locks are observed for all the proteins in our large conformational change set (see Figures SI-5-16) with the exception GroEl, where the residues forming rigid pair upon protein closure seem to be more distributed over the protein structure. In this protein, pairs undergoing a large increase in their directional force constants upon closure involve residues from the apical and equatorial domains (which correspond respectively to the upper and lower parts of the cartoon representations in Figure 8). Remarkably, the conformational transition of GroEL was also investigated using normal modes by Uyar et al. in Ref. (Uyar et al., 2014), and turned out to be one of the most difficult case for predicting the protein closed conformation when starting from an open structure. These results suggest a complex conformational pathway between the two forms, which would result in the distribution of rigid residues pairs all over the protein structure. Furthermore, the GroEL monomer activity takes place within the large hom-oligomeric GroEL complex, where inter-subunit contacts, which cannot be addressed in this work, are likely to play an essential part in the protein function.

**Figure 8.**
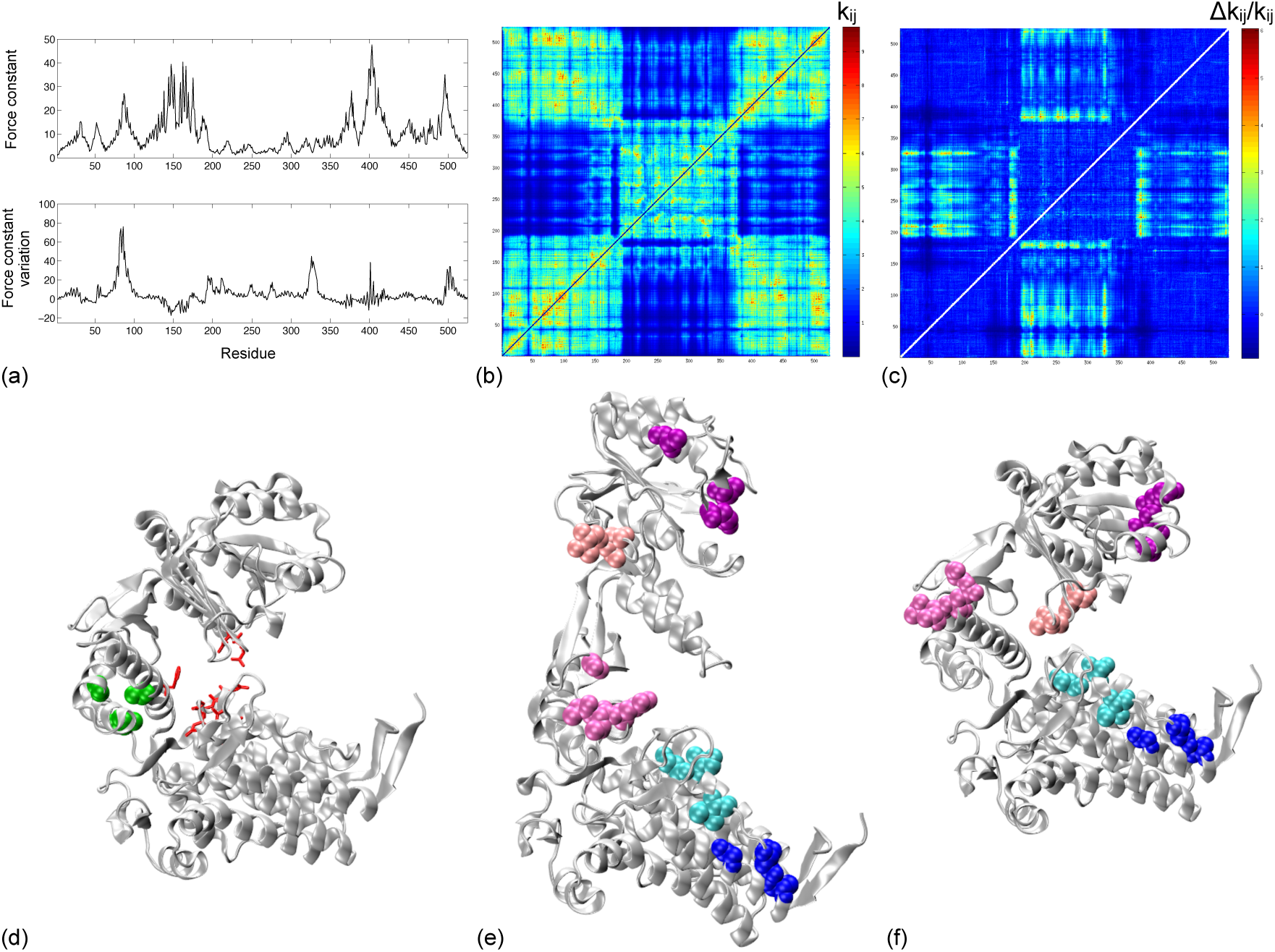
GroEL. (a) Upper panel: Rigidity profile for the protein in its open form. Lower panel: Force constant variations upon closure. (b) Directional force constants matrix for the protein in its open form. (c) Relative variation of the DFCs upon closure. (d) Cartoon representation of the protein in its closed form. Cartoon representations with the residues forming rigid pairs upon closure shown as van der Waals spheres: (e) Open structure (f) Closed structure.

**Coevolution analysis of the residues:** In our earlier work on guanylate kinase we noticed that the residues forming the structural lock seemed to occupy similar positions to the set of coevolved residues that were detected by Armenta-Medina et al. (Armenta-Medina et al., 2011) using statistical coupling analysis (SCA) in their work on adenylate kinase. This qualitative agreement also appears with the structural lock residues detected for AK in the present work, which are distributed on the LID, and NMP domains (see Figures SI-13e and f) and seem to overlap with the coevolved residues of Armenta-Medina et al. shown on Figure 1 in ref. (Armenta-Medina et al., 2011). In addition, coevolution has also been used to determine alternative conformational states in proteins by Sfriso et al. (Sfriso et al., 2016), or for the determination of protein domains (Granata et al., 2017). In order to get a more quantitative view of this phenomenon and to determine wether structural lock residues are in fact coevolved, we performed a systematic investigation of the residues co-evolution for 12 proteins presenting large conformational transitions in our dataset (the endonuclease homodimer was not included in this test) using the CoeViz (Baker and Porollo, 2016) online tool with the default parameters (multiple sequence alignement based on the UniProt UniRef90 base). The resulting coevolution (or conservation) scores for each residue pair were plotted as a function of the DFC variation for this pair upon closure. Figure 9 shows the resulting plots in the case of adenylate kinase. While coevolution and important DFC variations may target the same areas in the protein, Figure 9 shows there is no significant correlation between the two quantities independently of the metric used to compute the coevolution scores (**χ**^2^, MI, or Pearson correlation). Moreover, the different coevolution scores also correlate badly with each other. Similar results were obtained when testing alternative bases for the multiple sequence alignment (UniRef50, NCBI, NCBI90, NCBI70, and Pfam), and for all the tested proteins, as can be seen in Figure SI-17, so eventually, we cannot draw any final conclusions regarding the co-evolution of structural lock residues.

**Figure 9.**
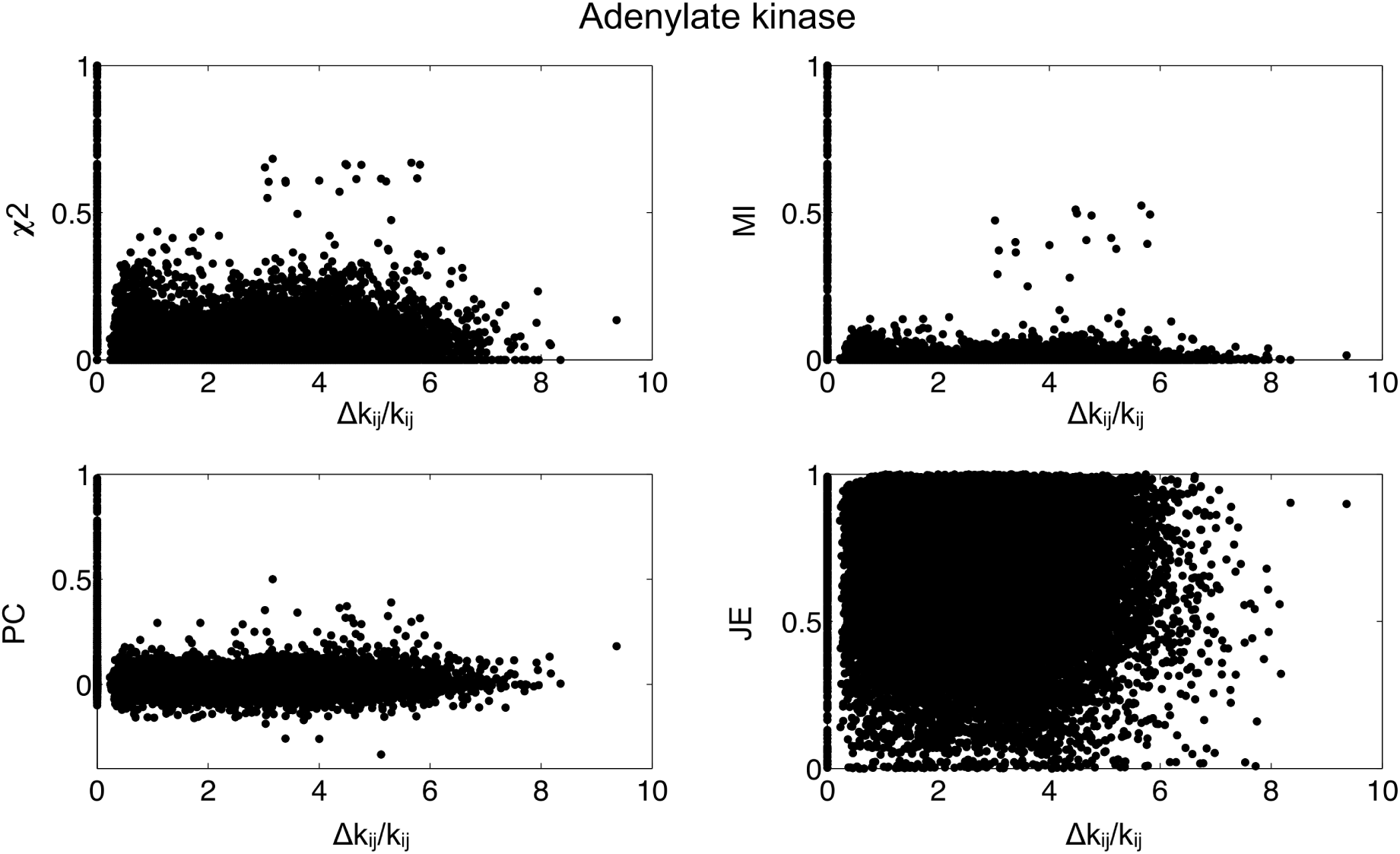
Distribution of coevolution scores from the CoeViz webserver (http://polyview.cchmc.org/) as a function of the relative variation in the DFCs upon closure for adenylate kinase.

## 4. Conclusion

We combine a coarse-grain elastic network protein representation and Brownian Dynamics simulations to investigate mechanical variations in proteins undergoing conformational changes, with a particular focus on proteins showing large scale motions with RMDSs between their open and closed forms larger than 4 Å. The variations in the residues force constants upon protein closure are highly heterogeneous along the sequence and difficult to predict. They are however tightly related to a protein’s biological activity, and residues undergoing the larger mechanical variation during the conformational changes are usually important for function. Interestingly, a protein’s average rigidity in its open form also provides a good estimate on wether it is likely to undergo large-scale motions or not.

For proteins presenting important conformational changes upon closure, we also calculated the directional force constants between all residue pairs. The resulting maps highlight how the proteins are structured in adjacent domains. Investigating the DFCs variation upon protein closure leads to the identification of a subset of residues whose side-chains are tightly interacting in the closed structure, and forming what we call a *structural lock*. Therefore, the calculation of DFCs brings additional information regarding a new class of functionally relevant residues in proteins with large-scale motions that are not directly involved in catalytic activity, but provide functional support by enhancing the closed structure stability. From a protein design perspective, the structural lock residues represent potential mutation targets to change the closed state stability in order to enhance or modulate protein function. On can also note that these pairs could not be identified using alternative methods such as rigidity profiles, since the individual residues do not correspond to rigid spots in the protein. Distance variation matrices, binary contact maps and normal mode analysis will highlight a much larger subset of residue pairs than the mechanical approach. Additional calculations of coevolution and conservation scores also failed to predict the specific residues that will form a structural lock within the protein closed conformation.

## Acknowledgments

This work was supported by the “Initiative d’Excellence” program from the French State (Grant “DYNAMO”, ANR-11-LABX-0011–01).

## Supporting Material

A table listing all proteins studied in this work and 17 additional figures are available as supporting information. In addition, the calculations results (rigidity files and directional force constant matrices) have been made available as an archive which can be downloaded here: https://owncloud.galaxy.ibpc.fr/owncloud/index.php/s/WYGTdxyuTunmxHw

